## Supplementary material for "Antibiotics promote intestinal growth of carbapenem-resistant *Enterobacteriaceae* by enriching nutrients and depleting microbial metabolites"

### SUPPLEMENTARY FIGURES

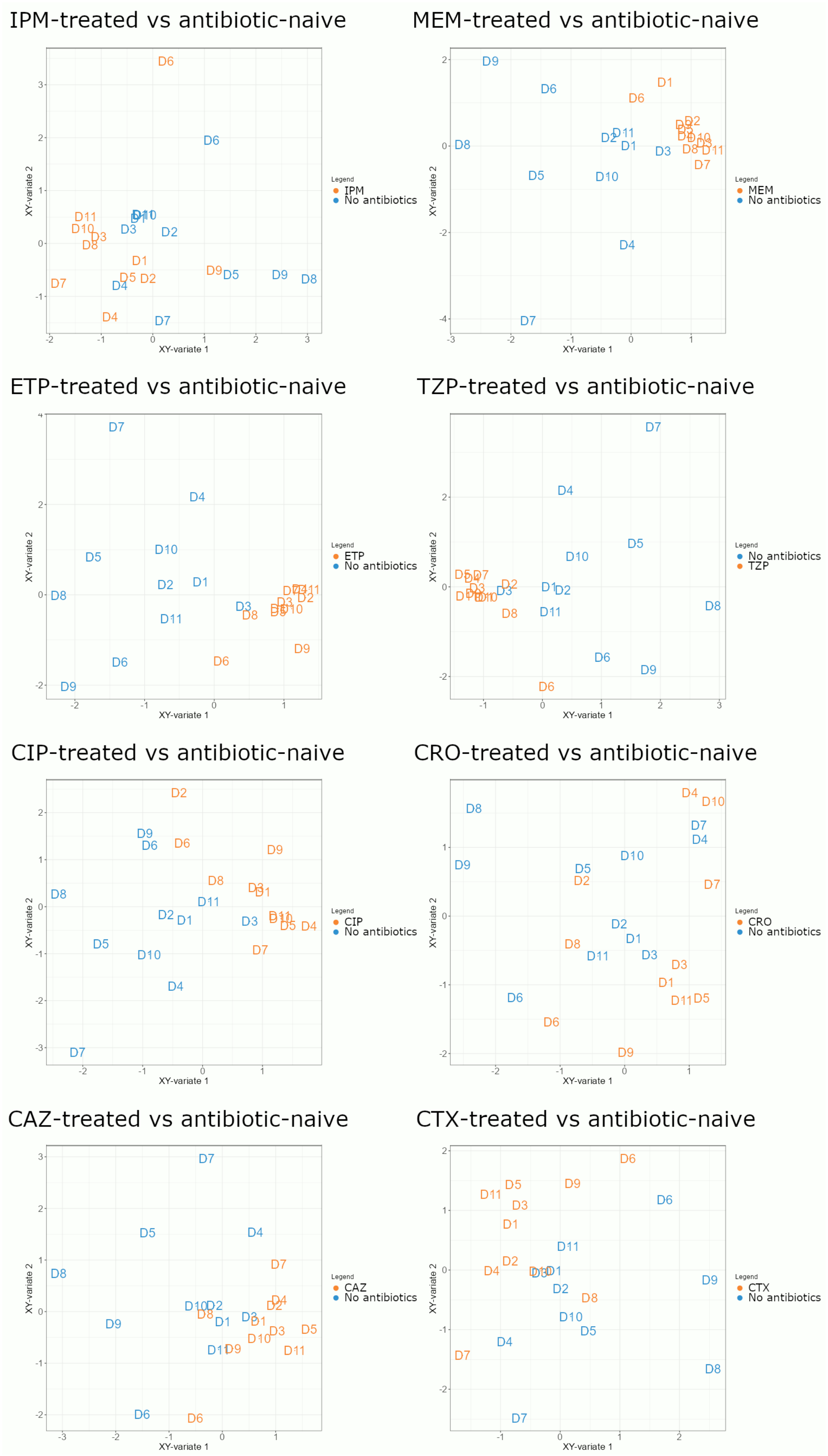


**Figure S1: Antibiotic treatment resulted in a separation of antibiotic-treated and antibiotic-naïve human faecal microbiota in rCCA representation unit plots.** rCCA models correlating 16S rRNA gene sequencing data (family level) and $^{1}$H-NMR spectroscopy data. Representation of units (aka samples) for the first 2 canonical variates showing the correlations between variables antibiotic-treated (orange) and antibiotic-naïve (blue). n = 11 healthy faecal donors, D1-D11 = samples from donor number 1-11.

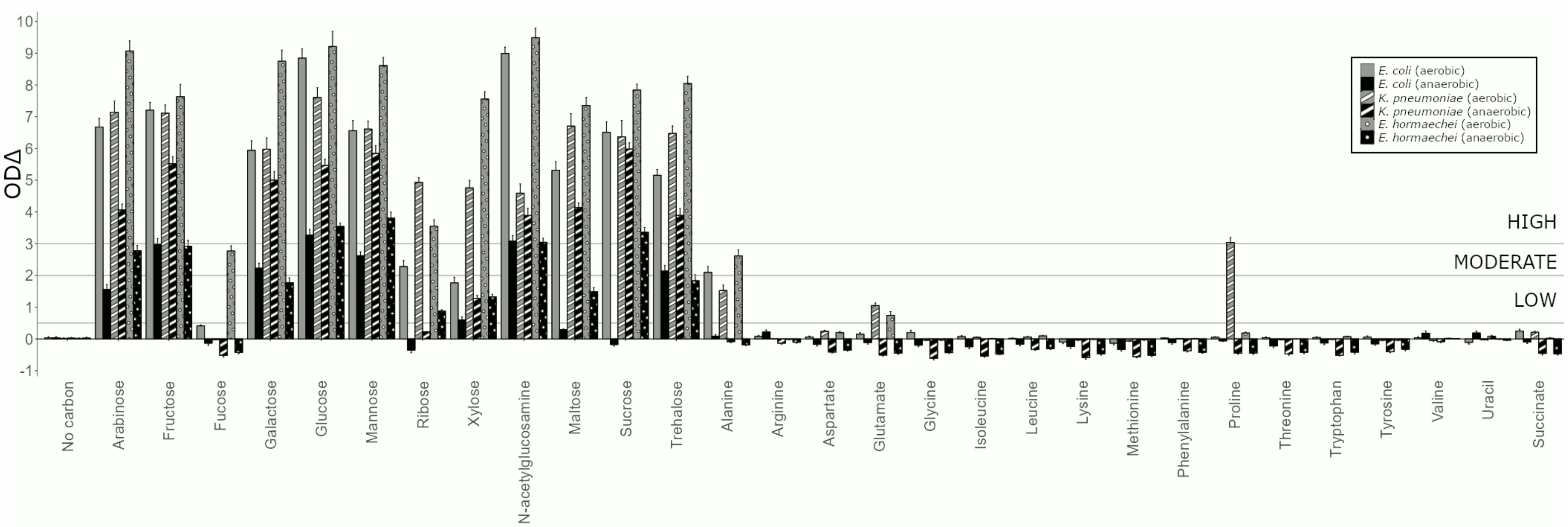


**Figure S2: Individual carbon sources support the growth of CRE at high, moderate, or low levels.** Functional differences (ODΔ) between growth on a carbon source versus growth on the no carbon control for *E. coli* ST617, *K. pneumoniae* ST1026, and *E. hormaechei* ST278 grown under anaerobic or aerobic conditions are summarised with the sum of functional differences (ODΔ) quantifying the magnitude of differences between two growth curves. ODΔ >3 indicated high growth, ODΔ between 2-3 indicated moderate growth, ODΔ between 0.5-2 indicated low growth, ODΔ <0.5 indicated negligible or no growth. Error bars indicate 95% confidence intervals. Growth of each isolate was measured with 6 replicates in 2-3 independent experiments.

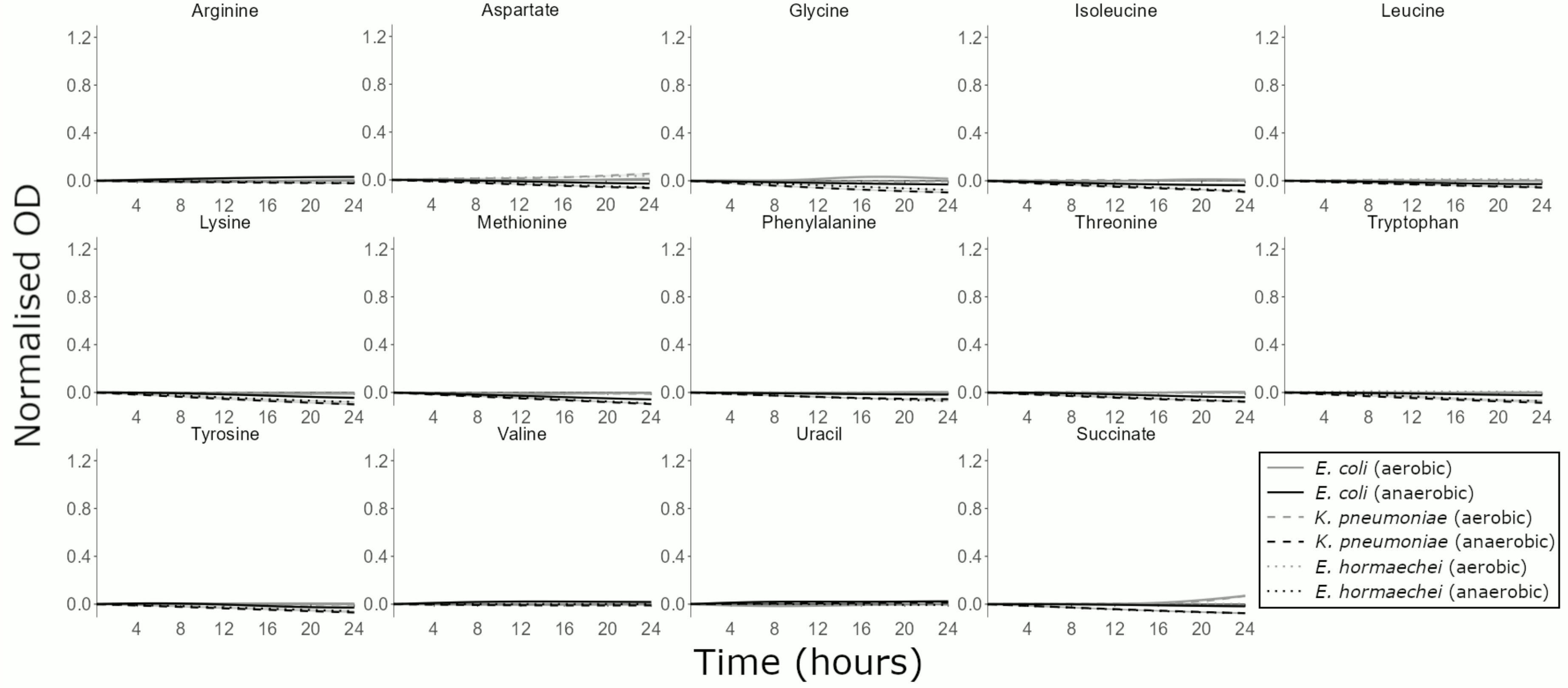


**Figure S3: CRE growth was not supported by some individual carbon sources that were elevated in antibiotic-treated faecal microbiota.** AMiGA-predicted growth curves for *E. coli* ST617, *K. pneumoniae* ST1026, and *E. hormaechei* ST278 grown on M9 minimal medium supplemented with a single carbon source (or water as the no carbon control) under anaerobic or aerobic conditions. Growth of each isolate was measured with 6 replicates in 2-3 independent experiments. The predicted mean of growth is shown with bold lines and the predicted 95% credible intervals are shown with the shaded bands.

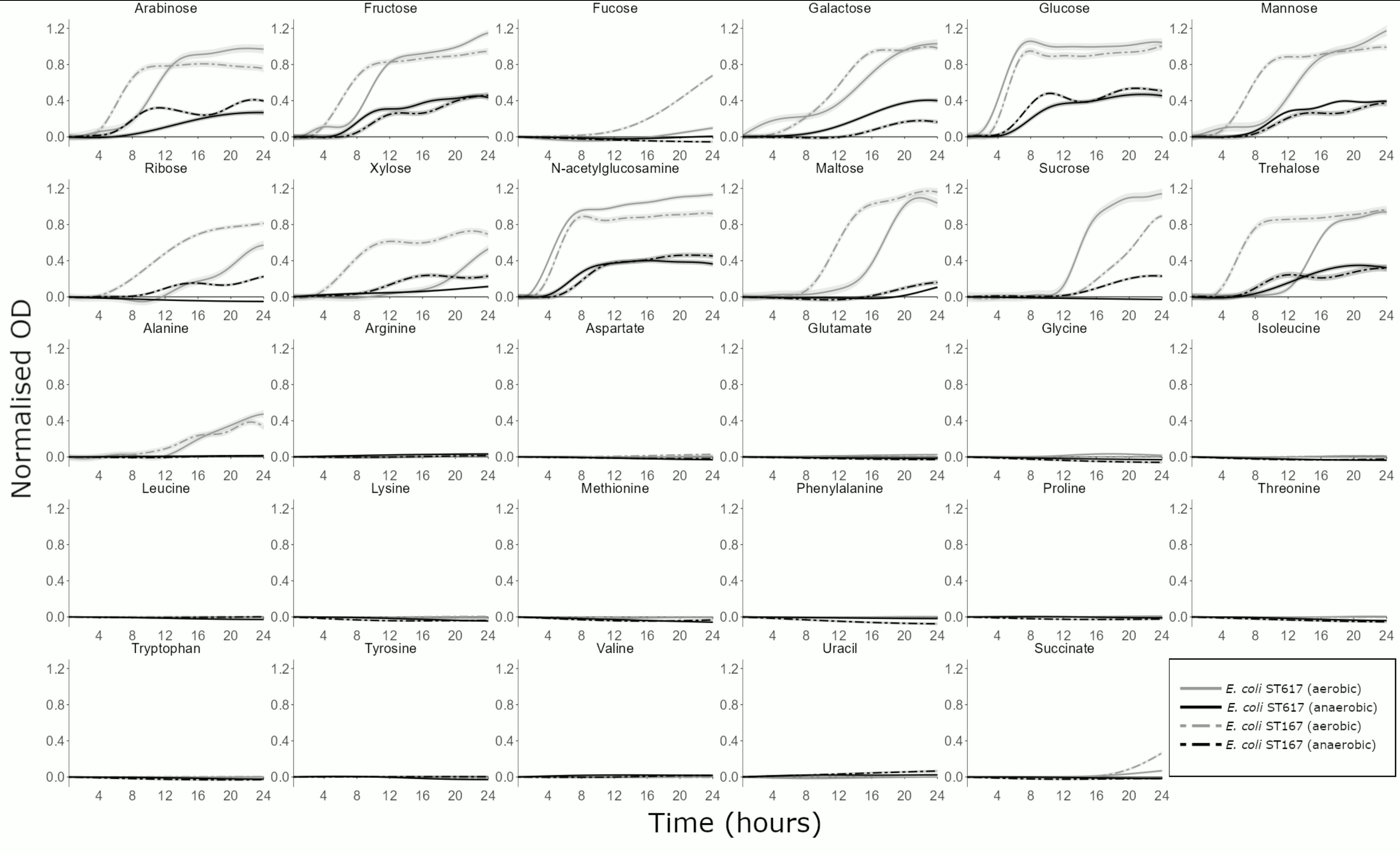


**Figure S4: Carbapenem-resistant *E. coli* growth was supported by many individual carbon sources that were elevated in antibiotic-treated faecal microbiota, but showed some between strain differences.** AMiGA-predicted growth curves for *E. coli* ST617 and *E. coli* ST167 grown on M9 minimal medium supplemented with a single carbon source (or water as the no carbon control) under anaerobic or aerobic conditions. Growth of each isolate was measured with 6 replicates in 2-3 independent experiments. The predicted mean of growth is shown with bold lines and the predicted 95% credible intervals are shown with the shaded bands.

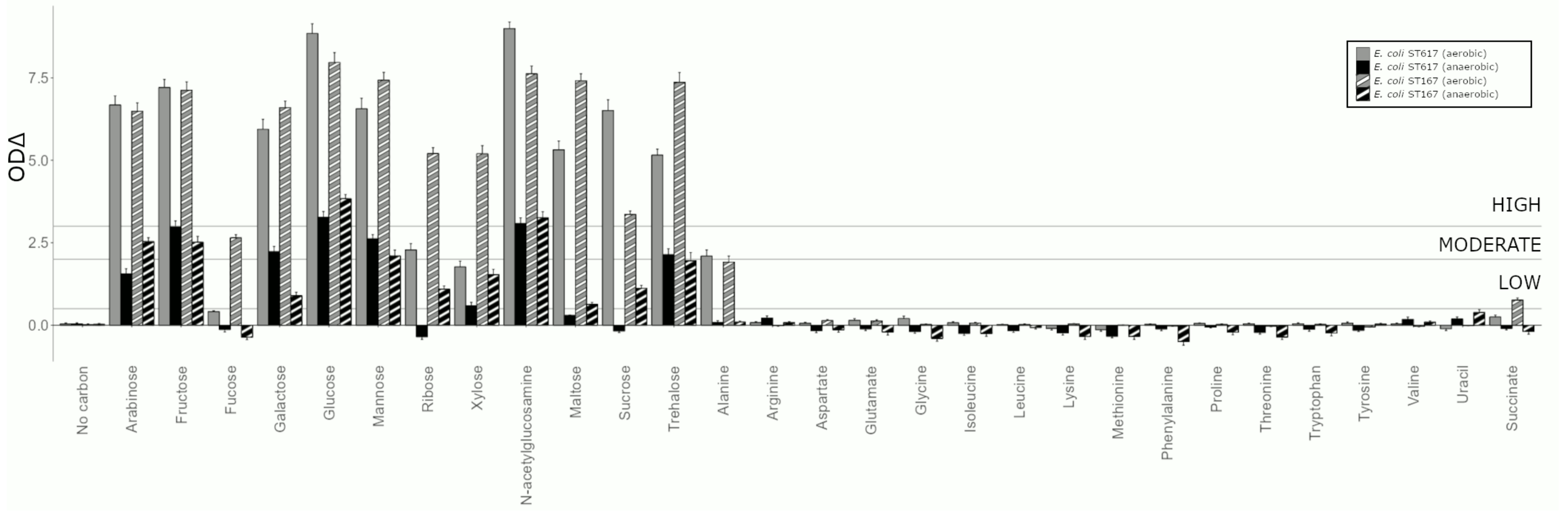


**Figure S5: Individual carbon sources support the growth of carbapenem-resistant *E. coli* at high, moderate, or low levels.** Functional differences (ODΔ) between growth on a carbon source versus growth on the no carbon control for *E. coli* ST617 and *E. coli* ST167 grown under anaerobic or aerobic conditions are summarised with the sum of functional differences (ODΔ) quantifying the magnitude of differences between two growth curves. ODΔ >3 indicated high growth, ODΔ between 2-3 indicated moderate growth, ODΔ between 0.5-2 indicated low growth, ODΔ <0.5 indicated negligible or no growth. Error bars indicate 95% confidence intervals. Growth of each isolate was measured with 6 replicates in 2-3 independent experiments.

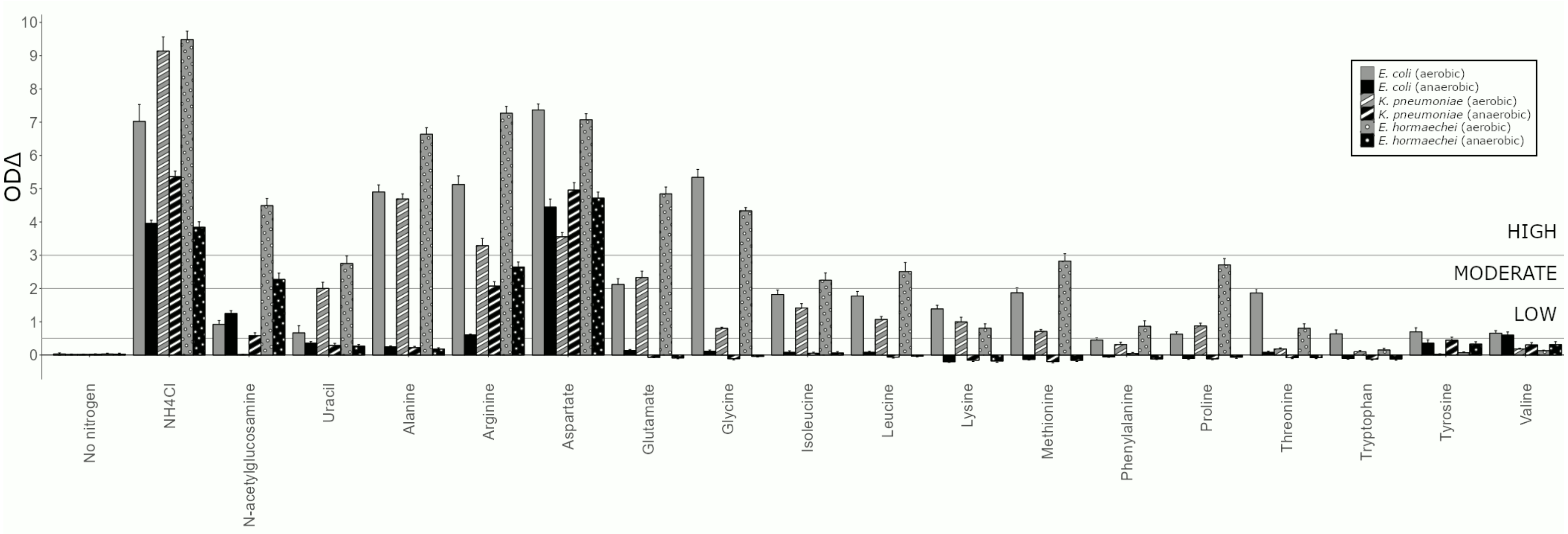


**Figure S6: Individual nitrogen sources support the growth of CRE at high, moderate, or low levels.** Functional differences between growth on a nitrogen source versus growth on the no nitrogen control for *E. coli* ST617, *K. pneumoniae* ST1026, and *E. hormaechei* ST278 grown under anaerobic or aerobic conditions are summarised with the sum of functional differences (ODΔ) quantifying the magnitude of differences between two growth curves. ODΔ >3 indicated high growth, ODΔ between 2-3 indicated moderate growth, ODΔ between 0.5-2 indicated low growth, ODΔ <0.5 indicated negligible or no growth. Error bars indicate 95% confidence intervals. Growth of each isolate was measured with 6 replicates in 2-3 independent experiments.

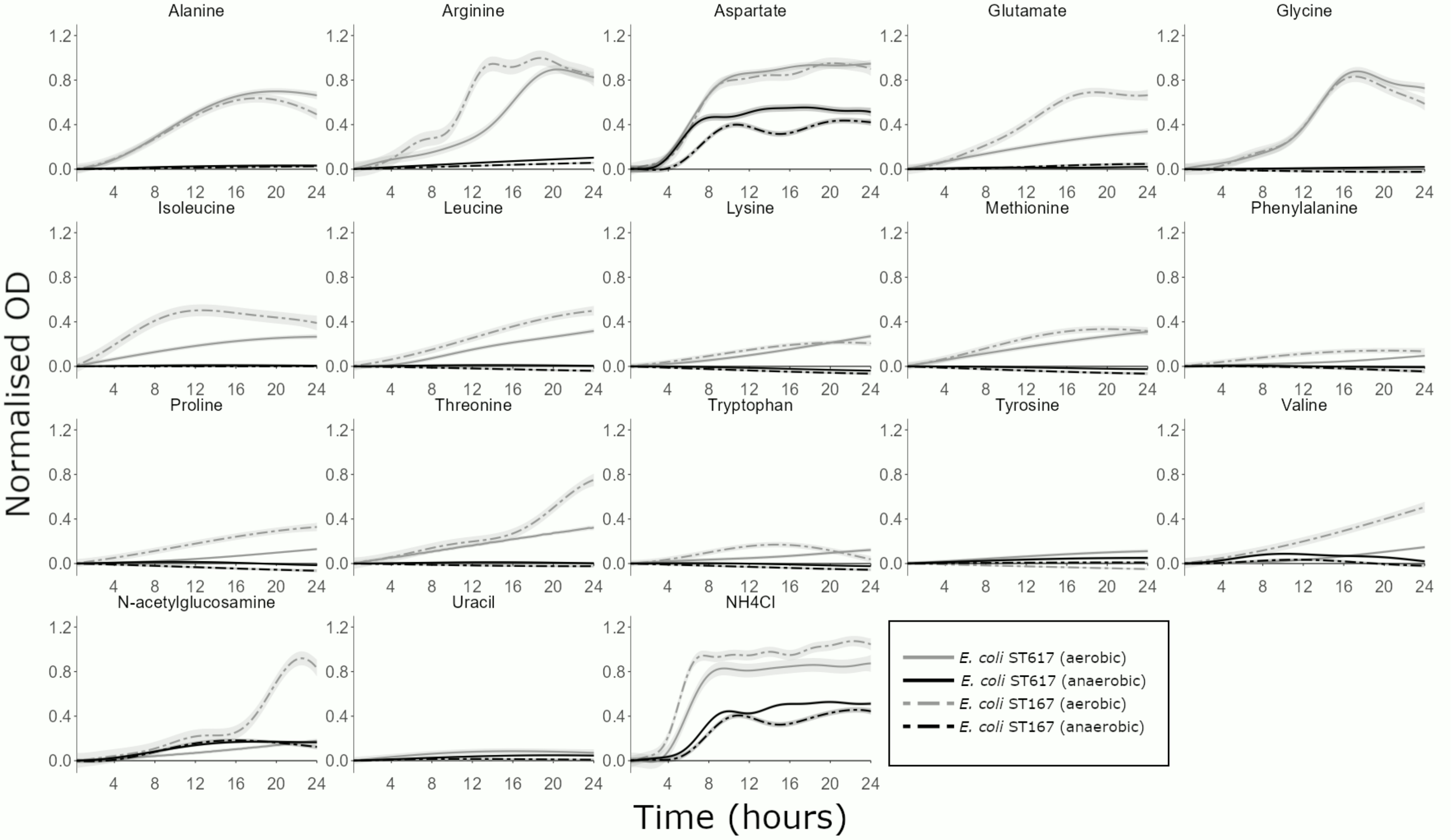


**Figure S7: Carbapenem-resistant *E. coli* growth was supported by individual nitrogen sources that were elevated in antibiotic-treated faecal microbiota, but showed some between strain differences.** AMiGA-predicted growth curves for *E. coli* ST617 and *E. coli* ST167 grown on M9 minimal medium supplemented with a single nitrogen source (or water as the no nitrogen control) under anaerobic or aerobic conditions. Growth of each isolate was measured with 6 replicates in 2-3 independent experiments. The predicted mean of growth is shown with bold lines and the predicted 95% credible intervals are shown with the shaded bands. $NH_4Cl$ was used as a positive control for growth as a sole nitrogen source.

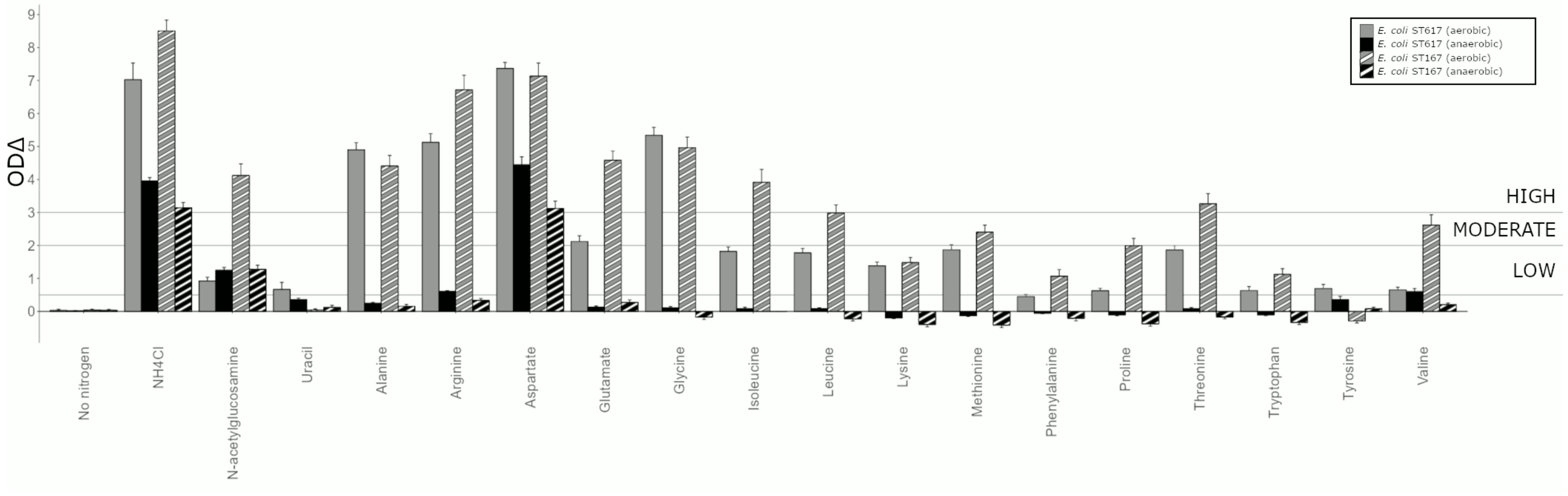


**Figure S8: Individual nitrogen sources support the growth of carbapenem-resistant *E. coli* at high, moderate, or low levels.** Functional differences between growth on a nitrogen source versus growth on the no nitrogen control for *E. coli* ST617 and *E. coli* ST167 grown under anaerobic or aerobic conditions are summarised with the sum of functional differences (ODΔ) quantifying the magnitude of differences between two growth curves. ODΔ >3 indicated high growth, ODΔ between 2-3 indicated moderate growth, ODΔ between 0.5-2 indicated low growth, ODΔ <0.5 indicated negligible or no growth. Error bars indicate 95% confidence intervals. Growth of each isolate was measured with 6 replicates in 2-3 independent experiments.

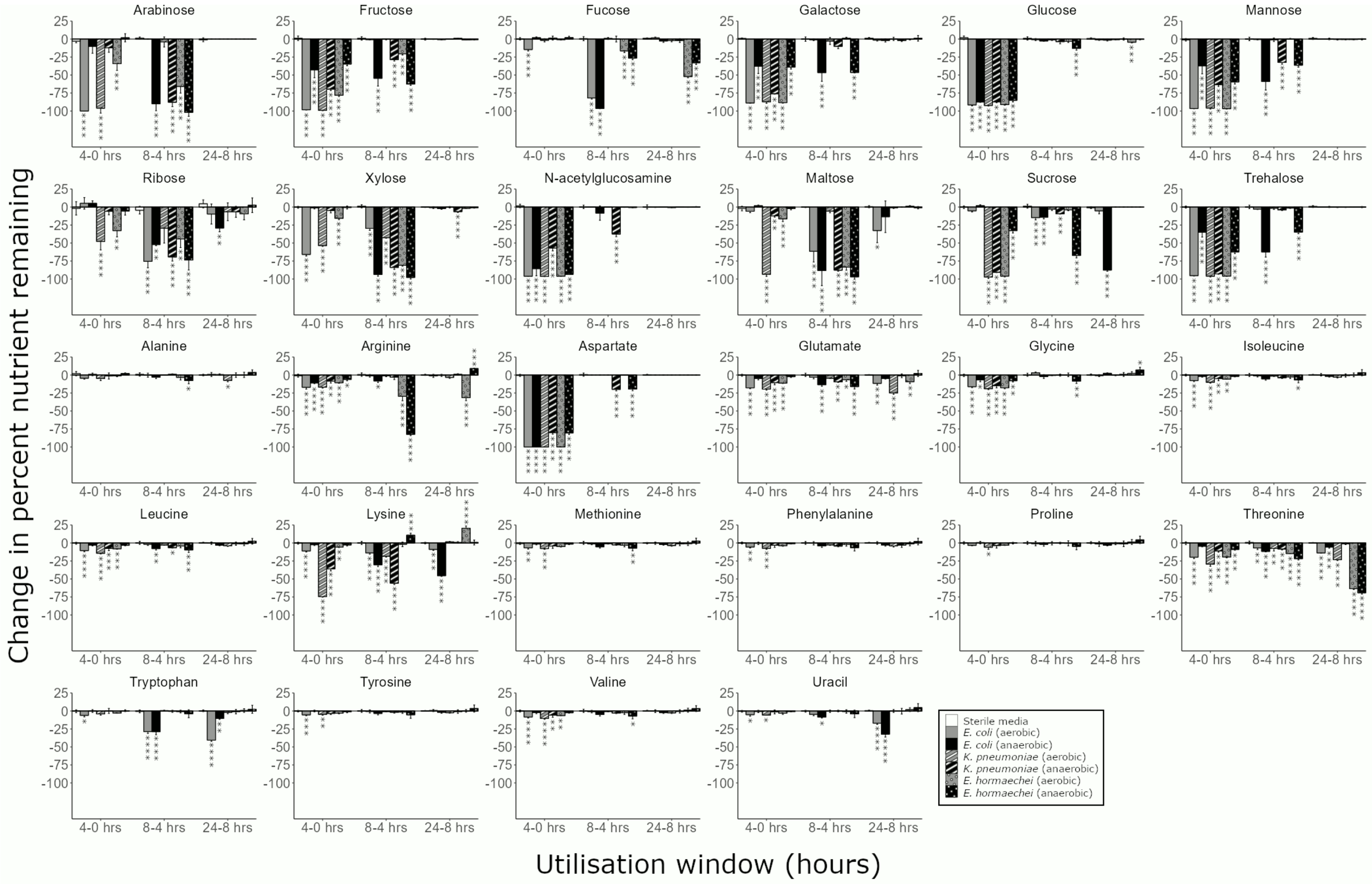


**Figure S9: Order of nutrient utilisation varies by CRE isolate and the presence or absence of oxygen.** Percent change in nutrients between subsequent time points by *E. coli* ST617, *K. pneumoniae* ST1026, or *E. hormaechei* ST278 grown under anaerobic or aerobic conditions in M9 minimal medium containing a mixture of 0.015% of each nutrient. Nutrient concentration was measured by $^1$H-NMR spectroscopy. Change in the percent of the nutrient remaining was calculated by subtracting the percent nutrient remaining at a time point from the percent nutrient remaining at the previous time point. Two-way mixed ANOVA followed by pairwise comparisons with Bonferroni correction. * = P ≤ 0.05, ** = P ≤ 0.01, *** = P ≤ 0.001, **** = P ≤ 0.0001, n=3 replicates for each CRE isolate and n=4 replicates for the sterile media controls.

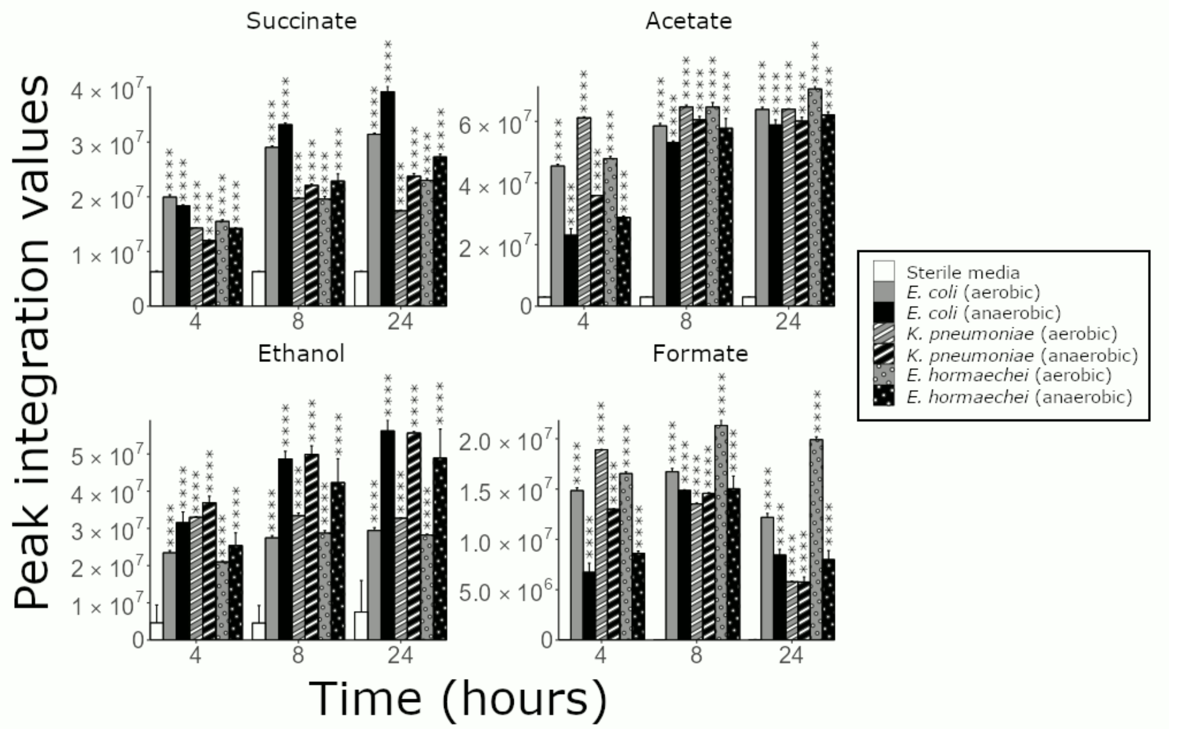


**Figure S10: CRE isolates produce metabolites following incubation with a mixture of nutrients.** Metabolite production by *E. coli* ST617, *K. pneumoniae* ST1026, or *E. hormaechei* ST278 grown under anaerobic or aerobic conditions in a minimal medium containing a mixture of 0.015% of each nutrient. Metabolite production was measured by $^{1}$H-NMR spectroscopy. The metabolite concentration was measured by the integration of a representative peak in the NMR spectrum after incubation for 4hrs, 8 hrs, and 24 hrs. Two-way mixed ANOVA followed by pairwise comparisons with Bonferroni correction. * = P ≤ 0.05, ** = P ≤ 0.01, *** = P ≤ 0.001, **** = P ≤ 0.0001, n=3 replicates for each CRE isolate.

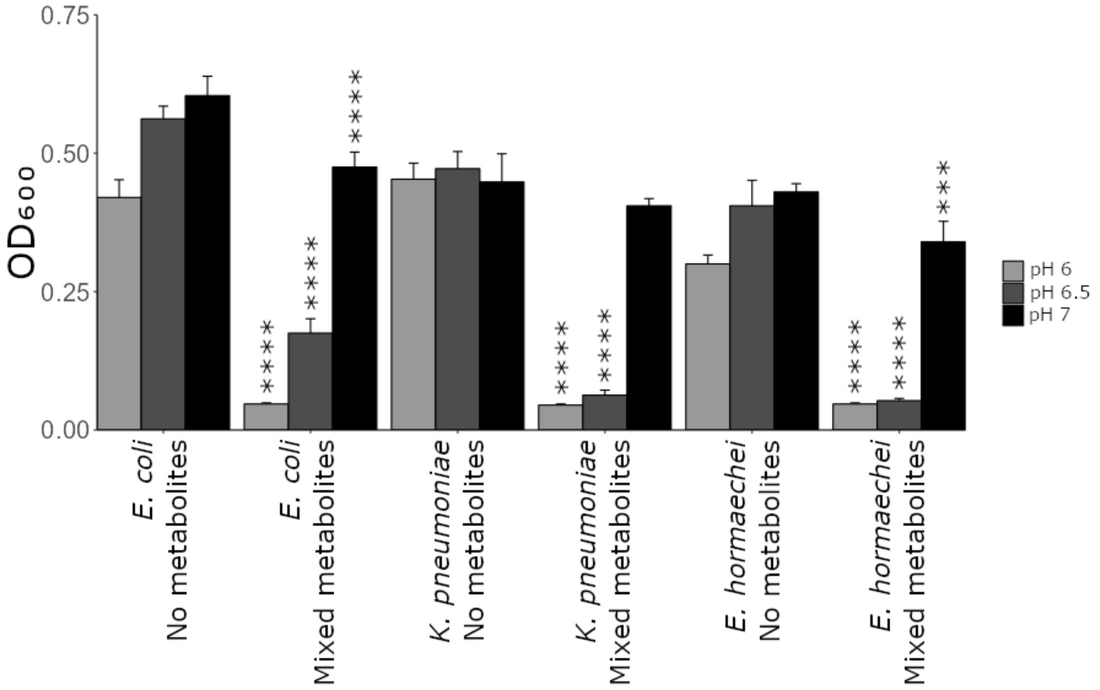


**Figure S11: CRE growth was inhibited by a mixture of microbial metabolites that were decreased in antibiotic-treated faecal microbiota when grown in a minimal medium supplemented with nutrients that were increased with antibiotics.** *E. coli* ST617, *K. pneumoniae* ST1026, or *E. hormaechei* ST278 were grown in M9 minimal medium (with a mixture of nutrients that were increased with antibiotics) supplemented with a metabolite mixture (mimicking average human faecal concentrations) or unsupplemented ("no metabolite" control) at pH 6, 6.5, or 7. Growth of each isolate was measured with 6 replicates in 2 independent experiments. Unpaired t-test comparing metabolite mixture to the no metabolite control at the corresponding pH. *** = $P \leq 0.001$, **** = $P \leq 0.0001$.

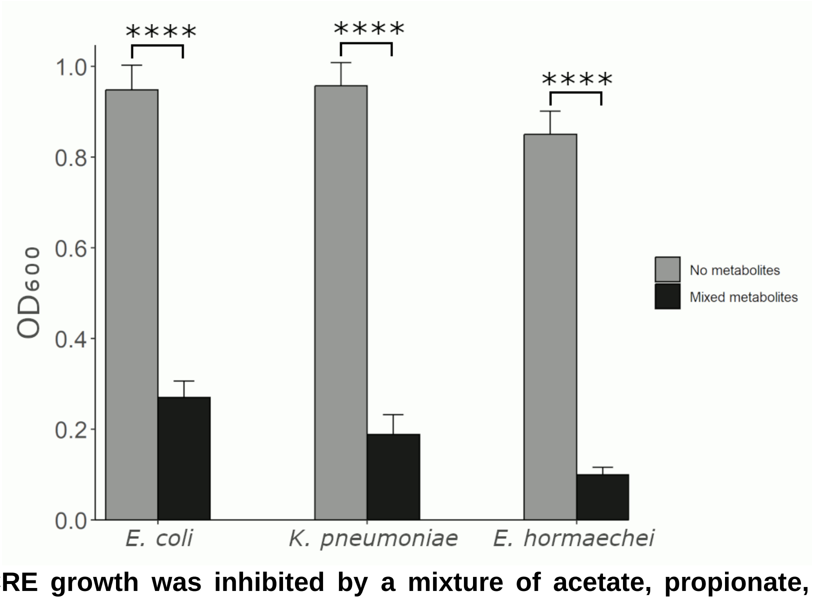


**Figure S12: CRE growth was inhibited by a mixture of acetate, propionate, butyrate, and valerate.** CRE growth was inhibited by a mixture of acetate, propionate, butyrate, and valerate at concentrations mimicking the average concentrations measured in human faeces. *E. coli* ST617, *K. pneumoniae* ST1026, or *E. hormaechei* ST278 were grown in LB supplemented with the metabolite mixture or unsupplemented ("no metabolite" control) at pH 6.5. Growth of each isolate was measured with 6 replicates in 2 independent experiments. Unpaired t-test comparing metabolite mixture to the no metabolite control. **** = $P \leq 0.0001$.

### SUPPLEMENTARY TABLES

**Table S1: Concentration of metabolites measured in human faeces from 12 healthy donors.**

| Metabolite | Lowest concentration (mM) | Average concentration (mM) | Highest concentration (mM) |
|---|---|---|---|
| Formate | 0.07 | 0.11 | 0.17 |
| Acetate | 10.54 | 64.08 | 122.73 |
| Propionate | 5.68 | 16.10 | 34.69 |
| Butyrate | 2.94 | 16.38 | 38.75 |
| Valerate | 0.52 | 3.67 | 12.01 |
| Isobutyrate | 0.33 | 2.20 | 5.28 |
| Isovalerate | 0.48 | 2.00 | 4.55 |
| Lactate | 0.26 | 0.67 | 1.20 |
| 5-aminovalerate | 0.29 | 1.06 | 4.15 |
| Ethanol | 0.64 | 10.46 | 59.62 |

**Table S2: Concentration of metabolites tested in the metabolite inhibition assays, based on measurements from human faecal samples from 12 healthy donors (found in Table S1).**

| Metabolite | Lowest concentration (mM) | Average concentration (mM) | Highest concentration (mM) |
|---|---|---|---|
| Formate | 0.05 | 0.10 | 0.15 |
| Acetate | 10 | 65 | 120 |
| Propionate | 5 | 15 | 35 |
| Butyrate | 3 | 15 | 40 |
| Valerate | 0.5 | 3.5 | 12 |
| Isobutyrate | 0.5 | 2 | 5 |
| Isovalerate | 0.5 | 2 | 5 |
| Lactate | 0.3 | 0.7 | 1.2 |
| 5-aminovalerate | 0.3 | 1 | 4 |
| Ethanol | 0.6 | 10 | 60 |

**Table S3: Whole genome sequencing results and antimicrobial resistance genes detected from draft genomes of CRE patient isolates.** See supplementary excel file.

**Table S4: MICs measured for CRE strains used in this study.**

| Strain | Meropenem, mg/L | Imipenem, mg/L | Ertapenem, mg/L | Piperacillin/tazobactam, mg/L |
|---|---|---|---|---|
| *E. coli* ST617 | 64 (resistant) | 16 (resistant) | 128 (resistant) | >128 (resistant) |
| *K. pneumoniae* ST1026 | 8 (intermediate) | 4 (intermediate) | 16-32 (resistant) | >128 (resistant) |
| *E. hormaechei* ST278 | 16 (resistant) | 4-8 (intermediate/resistant) | 32 (resistant) | >128 (resistant) |
| *E. coli* ST167 | 64 (resistant) | 16-32 (resistant) | 64-128 (resistant) | >128 (resistant) |
| *E. coli* ST410 | 2 (sensitive) | 2-4 (sensitive/intermediate) | 8 (resistant) | >128 (resistant) |
| *K. pneumoniae* ST258 | >128 (resistant) | 128 (resistant) | >128 (resistant) | >128 (resistant) |
| *K. pneumoniae* ST11 | 128 (resistant) | 128 (resistant) | >128 (resistant) | >128 (resistant) |

### SUPPLEMENTARY DISCUSSION

In this study we tested antibiotics that are frequently used clinically and known to promote the intestinal colonisation with CRE: carbapenems (MEM, IPM, ETP), penicillin/β-lactamase inhibitor (TZP), fluoroquinolones (CIP), and cephalosporins (CRO, CAZ, CTX)[1]. Although we have not tested all possible antibiotics that may promote the intestinal colonisation with CRE, these were a feasible number of antibiotics to test and logical choices to prioritise for this study. Faecal cultures treated with these 8 antibiotics caused broadly consistent enrichments of nutrients and depletions of metabolites (as shown in **Fig. 1b**). Future studies could explore the impact of other antibiotics on nutrient enrichment, metabolite depletion, and CRE growth.

Members of the *Bifidobacteriaceae*, *Bacteroidales*, and *Coriobacteriaceae* have been shown to utilise many of the nutrients that were elevated with antibiotics. *Bifidobacterium* have been shown to utilise most of the monosaccharides and disaccharides tested in this study[2]. Amino acid utilisation has not been well characterised in *Bifidobacterium*, however some *Bifidobacterium* species have been shown to utilise alanine, aspartate, glutamate, and threonine as sole carbon sources[3,4]. *Bacteroides* have been shown to utilise all the monosaccharides tested in this study and can utilise amino acids[5-7]. *Collinsella* can utilise most monosaccharides and disaccharides tested in this study and are known to metabolise amino acids[8,9].

We demonstrated that nutrient utilisation was influenced by the presence of other nutrients. Other studies have also shown that the presence of some nutrients influences the utilisation of others. Fucose has been shown to stimulate the utilisation of ribose by commensal *E. coli*[10]. Maltose utilisation gene expression was elevated in commensal *E. coli* grown on mucus[11]. Therefore, these results highlight the importance of studying nutrient utilisation as a mixture of nutrients in addition to their utilisation as sole nutrient sources.

A limitation of this study is that faecal microbiota were only analysed using $^1$H-NMR spectroscopy. Although this is an untargeted metabolic profiling technique that can measure a wide variety of relevant nutrients and metabolites of interest, it is not an exhaustive technique and cannot profile all nutrients and metabolites in our samples. There may be changes in other nutrients and metabolites that we did not measure (e.g. decreased primary fermentation of polysaccharides, mucin, or proteins with antibiotics). Therefore, if the utilisation of complex substrates by primary fermenters (such as *Bacteroides*) decreases, these substrates may provide an additional nutrient source for CRE. This potential role for primary fermenters also highlights the importance of continued investigation of the role of other nutrients and metabolites in promoting or inhibiting CRE intestinal colonisation and expansion.

We used faecal samples from 12 healthy human donors to quantify the average, minimum, and maximum metabolite concentrations to test in the experiments outlined in **Fig. 8**. Although this is not a large cohort of healthy human donors, the metabolite concentrations measured from faeces in our study were comparable to concentrations measured in other studies (where sample sizes ranged from 5-93 faecal donors)[12,13]. Future studies should measure the concentration of these 10 metabolites in a larger cohort of healthy human donors to confirm the metabolite concentrations found in healthy human faeces.

There were some limitations associated with the mouse experiments outlined in this study. Firstly, it is important to remember that mice are not a perfect model for humans for microbiome studies. There are differences in the microbiome taxonomic compositions and diets of humans and mice, among other factors[14]. Although we do not see the exact same changes in nutrients and metabolites in human and mouse faeces in response to TZP, our data demonstrates the same overall trends with TZP treatment: an increase in monosaccharides, disaccharides, and amino acids, and a decrease in SCFAs, BCFAs, ethanol, and lactate. Next, in this study mice were housed 5 per cage rather than individually. As mice engage in coprophagia, it is possible that mice were reinoculated with carbapenem-resistant *E. coli* that were ingested from faeces[15]. This may be improved by housing mice individually, however mice also ingest their own faecal pellets and so a certain degree of coprophagy is difficult to avoid in mouse experiments. As the mice caged together are from the same treatment group, faeces from all mice in the same cage are likely similar to each other. Therefore, we believe ingested faeces would be comparable in individually housed mice and group housed mice. Finally, although our mouse studies were designed using the appropriate power calculations, they were performed as one independent experiment. Designing an experiment to achieve a desired power (e.g. $\alpha = 0.05$ and a power of 0.80) and repeating this same experimental design several times may result in a higher power than is needed and overpower the study, resulting in the use of more animals than is needed[16]. However, it is important to replicate studies to ensure the results are reproducible. Future studies that continue the development of this metabolite mixture as a new treatment should repeat this experiment while also generating new data to make the most of this repetition – e.g. testing multiple doses of the metabolite mixture[17,18].

We did not measure the concentration of the supplemented metabolites in the mouse faeces or intestinal content after the metabolite mixture was administered to mice. However, SCFAs are routinely administered as their triglyceride versions as it is well established that these compounds are effective at increasing the luminal concentrations of the SCFAs in the distal gut[18]. Previous studies that administered glycerol tributyrate (aka tributyrin) to antibiotic-treated mice showed a significant increase in butyrate concentration in the colon contents and caecal contents of mice, where CRE are known to colonise[17,18]. Future studies should quantify metabolite concentrations in mouse faeces and in different segments of the gastrointestinal tract following administration of the metabolite mixture to better define the pharmacokinetics of this potential new therapeutic.
